# Cyclone: an accessible pipeline to analyze, evaluate and optimize multiparametric cytometry data

**DOI:** 10.1101/2023.03.08.531782

**Authors:** Ravi K. Patel, Rebecca G. Jaszczak, Im Kwok, Nicholas D. Carey, Tristan Courau, Daniel Bunis, Bushra Samad, Lia Avanesyan, Nayvin W. Chew, Sarah Stenske, Jillian M. Jespersen, Jean Publicover, Austin Edwards, Mohammad Naser, Arjun A. Rao, Leonard Lupin-Jimenez, Matthew F. Krummel, Stewart Cooper, Jody Baron, Alexis J. Combes, Gabriela K. Fragiadakis

**Affiliations:** University of California San Francisco (UCSF) CoLabs, UCSF, San Francisco, CA 94143, USA; Department of Pathology and ImmunoX Initiative, UCSF, San Francisco, CA 94143, USA; UCSF, San Francisco, CA 94143, USA; Department of Medicine, Division of Gastroenterology UCSF, San Francisco, CA 94143, USA; UCSF Liver Center, UCSF, San Francisco, CA 94143, USA; The Ibrahim El-Hefni Liver Biorepository (IELBC), San Francisco, CA 94115, USA; Division of General and Transplant Hepatology, California Pacific Medical Center Research Institute, San Francisco, CA 94115, USA; UCSF Immunoprofiler Initiative, San Francisco, CA 94143, USA; Department of Medicine, Division of Rheumatology, UCSF, San Francisco, CA 94143, USA

## Abstract

In the past decade, high-dimensional single cell technologies have revolutionized basic and translational immunology research and are now a key element of the toolbox used by scientists to study the immune system. However, analysis of the data generated by these approaches often requires clustering algorithms and dimensionality reduction representation which are computationally intense and difficult to evaluate and optimize. Here we present Cyclone, an analysis pipeline integrating dimensionality reduction, clustering, evaluation and optimization of clustering resolution, and downstream visualization tools facilitating the analysis of a wide range of cytometry data. We benchmarked and validated Cyclone on mass cytometry (CyTOF), full spectrum fluorescence-based cytometry, and multiplexed immunofluorescence (IF) in a variety of biological contexts, including infectious diseases and cancer. In each instance, Cyclone not only recapitulates gold standard immune cell identification, but also enables the unsupervised identification of lymphocytes and mononuclear phagocytes subsets that are associated with distinct biological features. Altogether, the Cyclone pipeline is a versatile and accessible pipeline for performing, optimizing, and evaluating clustering on variety of cytometry datasets which will further power immunology research and provide a scaffold for biological discovery.

## Introduction

The advent of high-dimensional single-cell technologies has transformed our ability to study the complex array of cell types, states, and behaviors comprising the immune system (1, 2). Single-cell proteomics including mass cytometry (CyTOF) and high-dimensional flow cytometry enable the detection of up to 50 extra- and intracellular proteins on hundreds of thousands of cells from a sample (3, 4). These technologies have been applied to a wide variety of patient cohorts and animal models to gain insights into immune set points, responses, and pathology including cancer, infection, hyperinflammatory disorders, and therapeutic intervention (5–8).

The high dimensionality of this data has necessitated the development and application of tools that can parse this data in a semi-automated way. This includes identifying cell populations via clustering algorithms (9–11), dimensionality reduction approaches for stratifying samples (12, 13), and visualization software and statistical packages for downstream analysis (14–16). Due to the diversity and complexity of the immune system, the use of clustering algorithms and dimensionality reduction has become increasingly standard for immune monitoring across tissues and species. However, to date a consensus process has not been established, rendering the comparison of this type of analysis difficult. While many tools have been both introduced and evaluated, many researchers such as wet-lab immunologists with limited computational experience struggle to navigate the vast landscape of tools for processing and analyzing cytometry datasets. Challenges in algorithm selection, run accessibility and scalability, and chaining the tools for each stage of analysis (preprocessing (17, 18), batch correction, clustering, and downstream analysis) pose significant barriers in the analysis of cytometry data. Moreover, many of these clustering algorithms require selecting a clustering resolution (i.e. selecting the number of clusters), which is largely arbitrary and may reduce the unsupervised nature of these methods. Therefore, an integrated cluster evaluation (19, 20) step is needed to compare different resolutions, guide clustering optimization, and facilitate a tool’s usage.

Here we present the <u>Cy</u>tometry <u>Cl</u>ustering <u>O</u>ptimization a<u>n</u>d <u>E</u>valuation (Cyclone) (<u>github.com/UCSF-DSCOLAB/cyclone/</u>) pipeline for the analysis of a wide range of cytometry data, including but not limited to mass cytometry (CyTOF), fluorescence-based cytometry, and multiplexed immunofluorescence (IF). Cyclone clusters data using FlowSOM (9) — selected based on scalability and fidelity to manual gating — and allows the users to optimize and evaluate cluster resolution based both on stability and user exploration. We present Cyclone’s performance on CyTOF datasets as well as other single-cell technologies including spectral flow cytometry and imaging. We designed Cyclone to be interoperable with outputs of the leading batch correction algorithms (21, 22) and to feed into accessible downstream analysis tools (16). We additionally accommodate both large datasets as well as showcase Cyclone’s performance on down-sampled data, increasing its accessibility to those with either extensive or limited computational resources. Leveraging Cyclone on a seven-color multiplexed immunofluorescence dataset obtained from human colorectal and kidney tumors, we identified a distinct tumor-associated classical dendritic cell subset. We share the R-based pipeline publicly along with extensive documentation for its use in conjunction with upstream and downstream tools. With this largely “plug and play” pipeline we hope to lower the barrier to entry into high-dimensional cytometry analysis and provide a consensus tool to the research community.

## Results

### Building a scalable pipeline for analysis of high dimensional cytometry data

To create a functional and accessible workflow for cytometry datasets, we were interested in building a pipeline optimized to meet criteria that address challenges in analyses of this nature (**Figure 1A**). This included 1) a framework optimized for interoperability, versatility, and ease of use, 2) designed to receive standard inputs from pre-processing steps including batch correction, and 3) provide a set of outputs that can easily serve as inputs to downstream analysis and visualization tools. The pipeline required both the selection of a scalable and accurate clustering algorithm robust to downsampling, as well as the ability to tune and evaluate clustering resolution to best meet the specifics of the biological system and scientific inquiry. Due to the wealth of existing single-cell technologies and the need of multiple measurement types to fully understand complex biological systems, we were additionally interested in developing a pipeline which would accommodate a range of single-cell cytometric data modalities including imaging.

**Figure 1.**
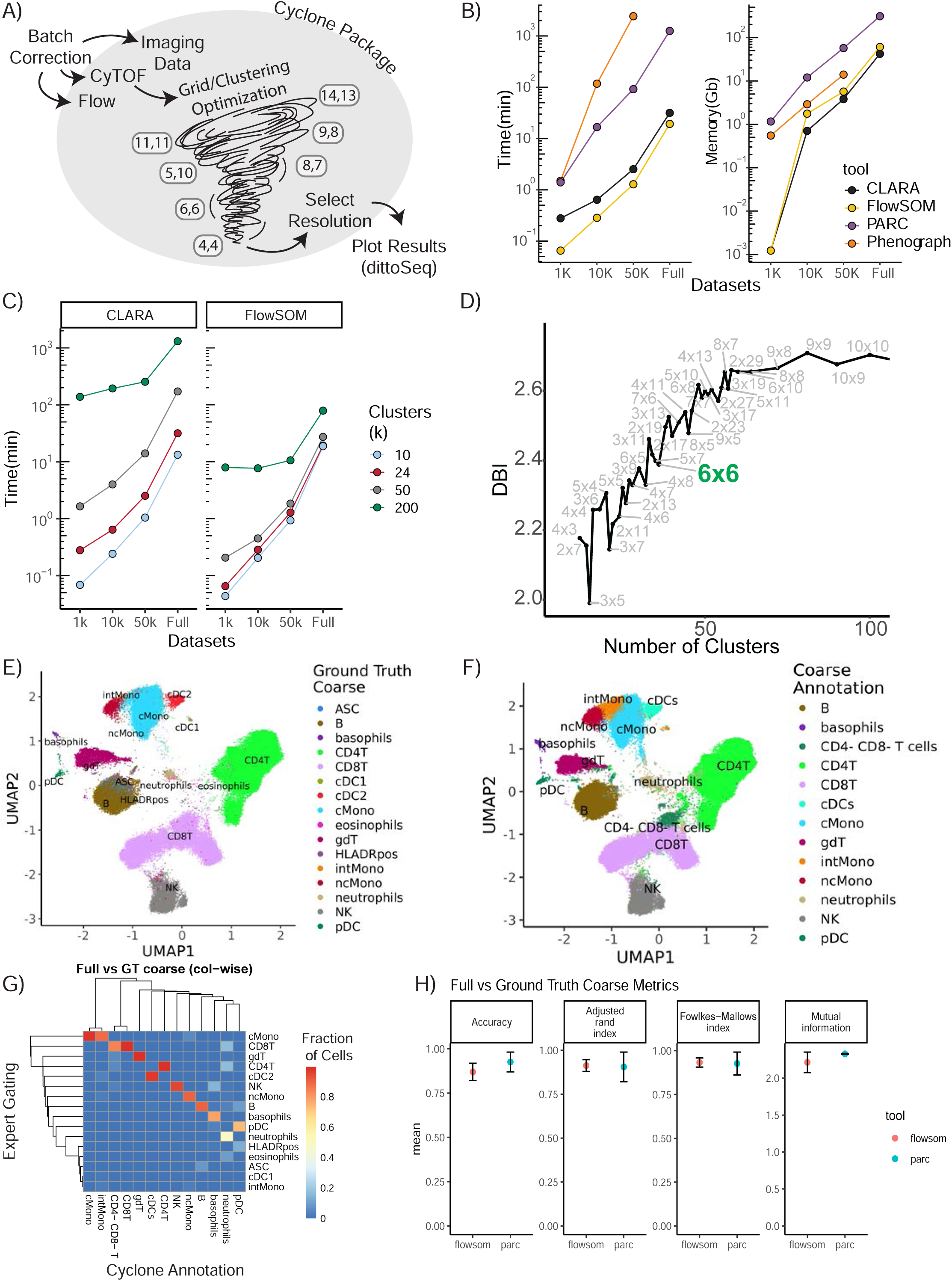
Building a scalable pipeline and optimizing clustering for analysis of high dimensional cytometry data. **A)** Graphical depiction of cyclone pipeline. Cyclone can intake multiple types of cytometry data, including Flow, CyTOF, and imaging data, and works for both raw and batch corrected data. FlowSOM clustering includes a grid/clustering optimization step, where a set list of grids are calculated. After the user selects a desired resolution, Cyclone generates useful plots as well as processed dataframes that can be easily handed off to other analysis tools and plotting packages, like dittoSeq or ggplot2. **B)** Clustering tool performance on data, through the lens of scalability (run time and memory required) of tested tools. CLARA and FlowSOM were similar in their time and memory requirements, while PARC and Phenograph have less feasible run time or memory requirements. **C)** Clustering time increases as cluster number (k) increases. FlowSOM still performs better than CLARA when measuring time to cluster, regardless of cluster number requested. **D)** Optimization of clustering via evaluation of the different resolutions, leveraging Davis-Bouldin index as indicator (subset of full number of grids assessed; full amount of grids evaluated in the supplement) **E)** Ground Truth expert “coarse”-level annotation identifying broad cell types based on manual gating. **F)** FlowSOM cluster annotation at “coarse”-level based on CyTOF panel expression. **G)** Heatmap comparing “coarse”-level annotations assignment based on the “ground truth” manual gating (rows) of full dataset vs FlowSOM cluster annotated by expert immunologist based on Cyclone cluster heatmap outputs (columns) **H)** Comparison metrics based on “coarse”-level annotations from two expert immunologists. Various performance metrics were used to assess the accuracy of clusters called in the FlowSOM clustering compared to ground truth.

A critical component of the pipeline was to select a clustering algorithm that could 1) scale to a large number of cells and parameters while maintaining reasonable run times and memory usage; and 2) recapitulate populations identified via expert manual gating. We therefore evaluated a subset of available clustering algorithms, focusing on popular algorithms shown to perform well in the literature. These included PhenoGraph (11), a graph-based community detection algorithm; CLARA (23), an extension of the partitioning around medoids algorithm; FlowSOM (9), which uses self-organizing map clustering; and PARC (10), a recent combinatorial graph-based algorithm optimized for scalability. We applied these algorithms to a CyTOF dataset measuring the expression of 42 proteins at the single-cell level on peripheral blood mononuclear cells (PBMC) from 17 individuals (Methods, **Supplemental Table 1**). We also subset the dataset to various sizes ranging from 1,000, 10,000, and 50,000 cells per individual to evaluate both speed and memory usage on a range of dataset sizes (**Figure 1B**). While memory usage was similar across algorithms (except for PhenoGraph performance on the full dataset, which did not conclude in a reasonable amount of time), FlowSOM and CLARA utilized the least runtime across dataset sizes. We therefore proceeded evaluating those two clustering algorithms on the full dataset. Notably, increasing requested cluster number identified by the algorithm across the variously sized datasets strongly increased runtime for CLARA as compared to FlowSOM (**Figure 1C**), which appeared more time efficient to cluster even larger cluster counts regardless of cell numbers. This indicated that CLARA takes more time than FlowSOM to perform a high-resolution clustering to identify cell subsets.

While our final pipeline does focus on R-based implementation for clustering (FlowSOM and CLARA) and permits maximum interoperability with other R based normalization, debarcoding, and batch correction packages, early investigations explored annotations from both R and python packages; thus, we benchmarked FlowSOM and PARC cluster annotations against manual gating (**Supplemental Figure 1**) to evaluate their performance in identifying cell populations. To compare the accuracy of annotating clusters based on CyTOF panel expression to gating Ground Truth, we developed a custom cell barcode scheme (See “Methods - FCS modifications”) to identify cells in the FCS and facilitate comparison of a cell’s Ground Truth gating-based assignment versus how the cell was annotated by two expert immunologists. To select a resolution/grid for annotation and evaluation, we calculated the Davies-Bouldin Index (DBI) (19), a within-dataset similarity metric used to evaluate cluster resolution, across a variety of cluster grid sizes (**Figure 1D, Supplemental Figure 2A**). Thirty-six clusters (6×6 grid, FlowSOM) or thirty-seven clusters (Resolution 1.3, PARC) were selected for independent annotation by two expert immunologists to compare to “ground truth” manual gates from a third expert immunologist. Both ground truth gating and cluster annotations were done for major immune cell populations (“coarse level” e.g. all CD4+ T cells subsets including Memory or Naïve are annotated as CD4+ T) (**Figures 1E, 1F**) and for more fine-grained sub-populations (“fine level”, **Supplemental Figure 2B-E**); these annotations were then visualized in UMAP space. Both FlowSOM and PARC coarse annotations performed well in the recapitulation of manual ground truth gating as captured by four evaluation metrics, including Accuracy, Adjusted rand index (24), Fowlkes-Mallows index (25), and Mutual information (26) (**Figure 1G, 1H**). As expected, while performance was not as strong on fine annotations, metrics showed reasonably accurate clustering for both clustering algorithms (**Supplemental Figure 2C**). Some sources of error included the fact that global clustering did not isolate small subsets including cDC1s or Antibody Secreting Cells (ASC) as their own cluster at the selected resolution due to its low abundance. In addition, the accuracy of some subsets of CD4+ and CD8+T cells varied due to their definition based on markers that have a continuum of expression rather than clear positive and negative expression (e.g. CD45RA). On the other hand, the unsupervised nature of clustering facilitate the identification of cell subset, such as B cell subsets based on the expression of CXCR5 expression or CD38 (**Figure 1G**). Notably, while the majority of clusters were easily recognizable and annotated, a small number of clusters were observed to be dispersed across the UMAP visualization (**Supplemental Figure 2F**), rather than having a more homogenous phenotype (**Supplemental Figure 2G**), which illuminated the utility of visualizing each cluster’s dispersion as an output for a developed pipeline. Taken together, these data demonstrate that while PARC and FlowSOM perform similarly compared to expert gating and have similar performance on low numbers of cells, FlowSOM outperforms when considering scalability to higher numbers of cells.

### The Cyclone pipeline as a method for data clustering and evaluation across resolutions for use in the analysis of cytometry data

Based on our observations regarding runtime, memory usage, cluster evaluation methods, and performance on manual gating recapitulation, we developed the Cyclone pipeline in R (**Figure 2**). We designed Cyclone to produce a series of outputs users can access to select cluster resolution, evaluate cluster quality, annotate clusters, and utilize resulting cluster statistics (phenotypes and abundance) for downstream analyses. To start Cyclone, the user provides FCS or matrix data files as well as files to specify cluster markers and file metadata. Cyclone expects any normalization and batch correction steps to be performed prior to use of the pipeline. For CyTOF data, this can be done by established R packages, including premessa (<u>github.com/ParkerICI/premessa</u>) for bead-based normalization and CytoNorm (21) or Cycombine (22) for batch correction. After reading in and arcsinh transforming the data, Cyclone calculates UMAP dimensionality reduction. Cyclone works with either FlowSOM or CLARA for clustering; selecting FlowSOM enables DBI-based cluster resolution optimization prior to user grid selection, while with CLARA the user selects a single resolution for clustering. If FlowSOM is selected, Cyclone then performs iterative optimization of clustering across a variety of cluster grid sizes which can then be compared using DBI. After user selection of desired grid, Cyclone performs clustering, generates summary matrices, proceeds through an optional SCAFFoLD step, and then generates output files and visualizations. While not providing batch correction, Cyclone does provide a means for assessing batch or any other input file metadata via UMAP (split by batch) visualization (**Figure 3A**) as well as clustered heatmaps of cluster frequency with batch or other metadata information overlaid as rugs (**Figure 3B**). In addition to a DBI plot for cluster resolution selection, Cyclone outputs UMAPs colored by cluster (**Figure 3C**) as well as heatmaps of marker expression per cluster are also generated for ease of cluster annotation (**Figure 3D**). Additional UMAPs and histogram plots are provided per cluster, showing the cluster’s distribution across the UMAP for an evaluation of cluster dispersion as an indication of cluster quality (**Figure 3E**). Feature UMAP plots of marker expression in all cells are also export by default (**Figure 3F**). Taken together, the Cyclone pipeline provides a means of clustering, as well as evaluating and annotating these clusters, to then be readily used in downstream analyses.

**Figure 2.**
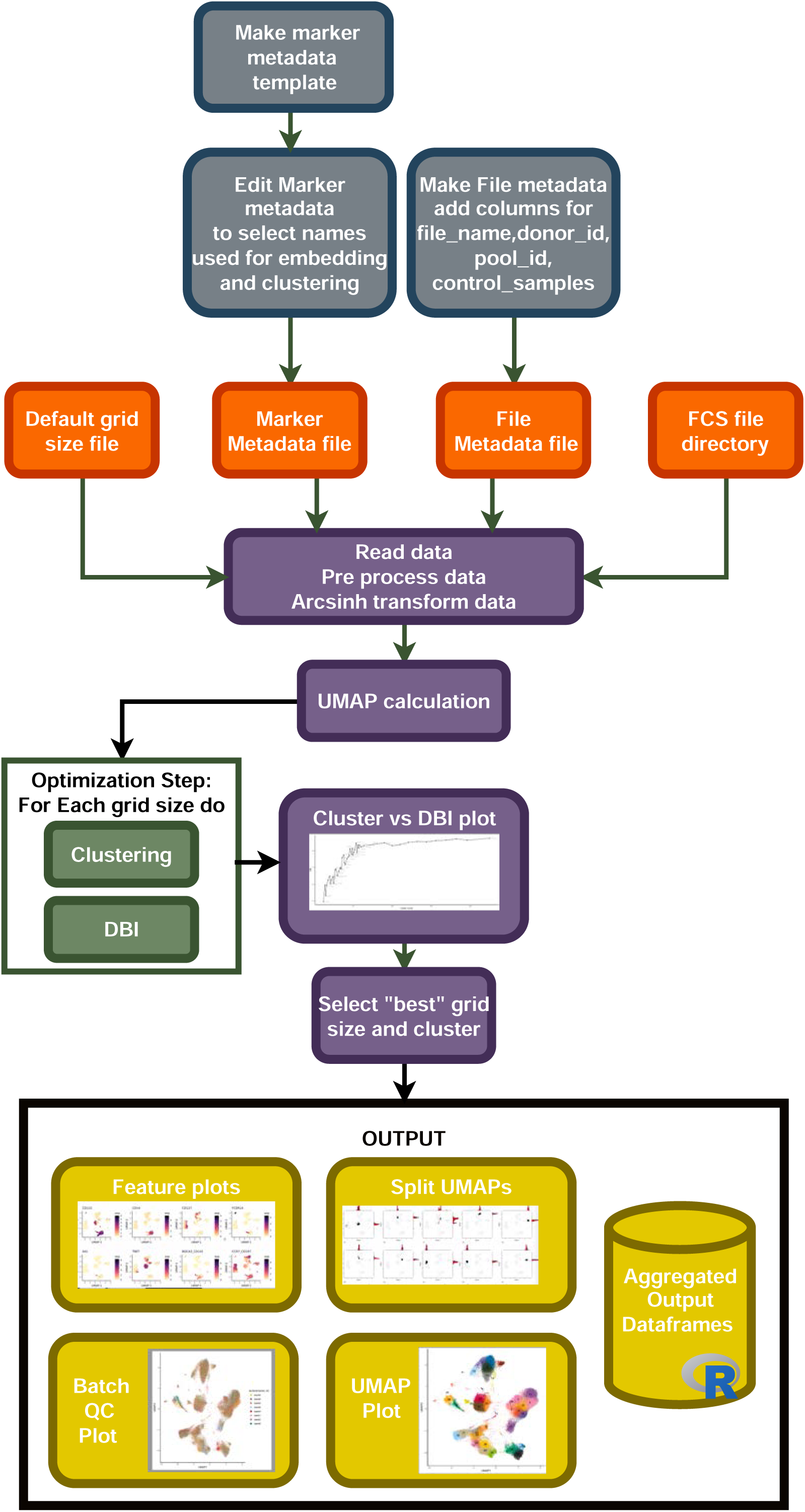
Integration of different pipeline pieces and packages into a single method: “Cyclone”. Steps of the Cyclone process. Using information from initial metadata files (see Supplemental Tables), FCS files are read into the pipeline. After arcsinh transformation, UMAP is calculated and clustering performed on a default range of grid sizes, resulting in a cluster VS DBI plot for clustering grid selection. After user input to select a specific grid, cyclone generates feature plots of each antibody included in the panel, split UMAPs to assess cluster dispersion, various QC heatmaps and UMAPs, and a final UMAP annotated by cluster number. Additionally, expression matrices and cell metadata matrices are saved.

**Figure 3.**
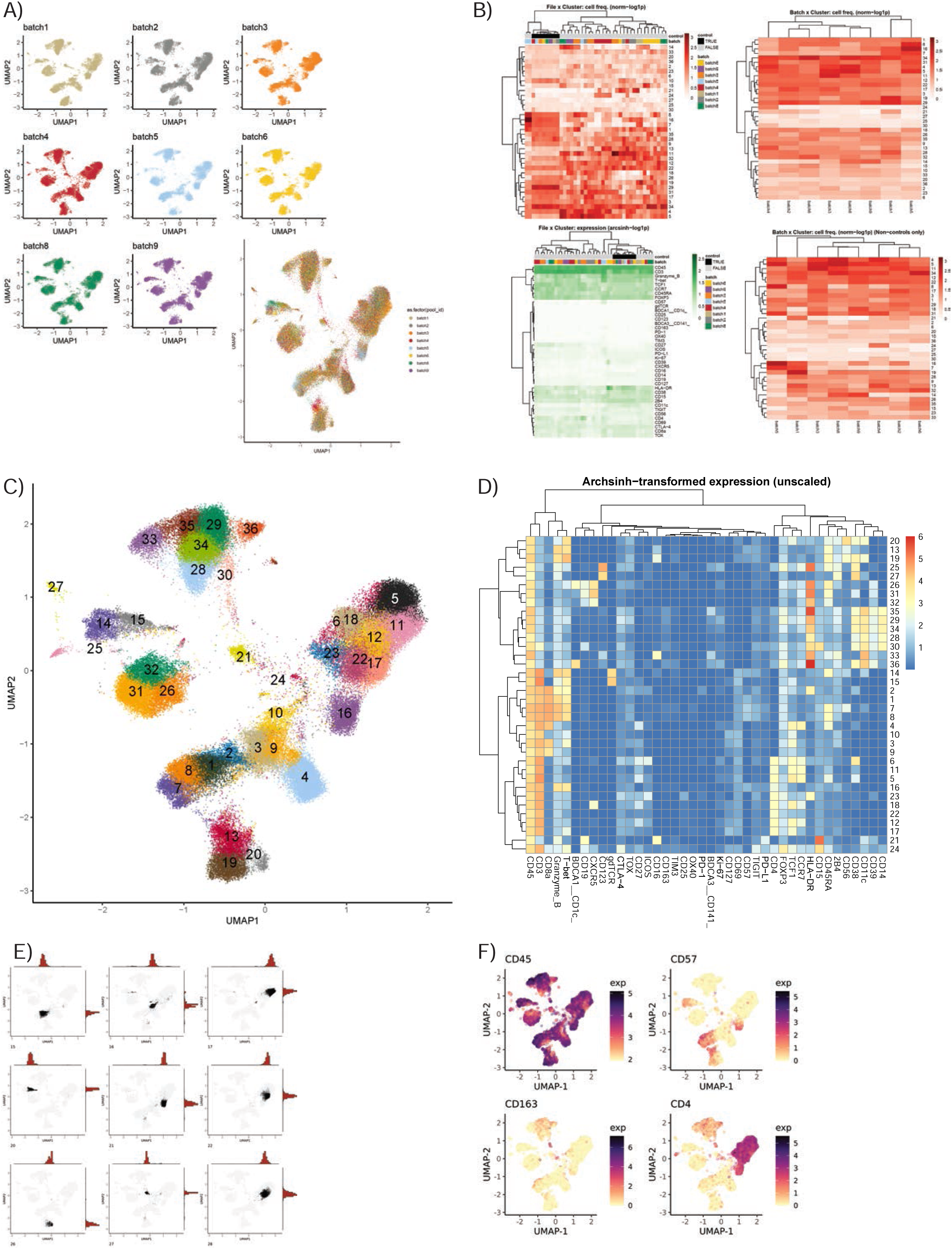
Evaluation and interpretation of default pipeline outputs and results. **A)** UMAPs of batch information. If batches were a part of the CyTOF run, Cyclone exports UMAP plots split by batch information to assess batch correction or batch effect. **B)** Heatmaps of file x cluster depicting various QC statistics such as cell frequency per cluster, batch per cluster, and arcsign transformed data. **C)** UMAP annotated by cluster number, based on user-selected grid. **D)** Heatmap of median archsinh-transformed expression (unscaled) per cluster, used to annotate clusters. **E)** Example plots of UMAP + histogram showcasing cluster density/dispersion. Example plots of protein feature expression. **F)** Example plots of protein feature expression.

### Cyclone provides accessibility and interoperability with upstream processing and downstream analyses

We developed Cyclone to be easily leveraged by researchers with various background, especially those with limited coding knowledge and minimal computational resources. We also prioritized interoperability with both upstream and downstream tools for cytometry data processing and analysis (**Figure 4A**). In addition to significant documentation (including vignettes to get users started with the pipeline), we evaluated Cyclone on downsampled datasets to determine whether downsampling to fewer cells could provide an alternative for users to run Cyclone locally rather than needing additional compute resources for analyses. Thus, we downsampled our evaluation dataset to 50,000 cells per sample and coarsely annotated the resulting optimized clusters (**Figure 4B, Supplemental Figure 3A, 3B**). We then compared the accuracy of these coarse downsampled annotations to the coarse annotations on the full dataset, and encouragingly found strong concordance between the full and downsampled datasets (**Figure 4C**). When doing fine annotations (**Supplemental Figure 3C**), metrics again showed reasonably accurate annotations when compared to the full dataset fine annotations (**Supplemental Figure 3D**). Thus, data downsampling presents a practical option for dataset clustering with Cyclone should compute resources be a challenge for some users.

**Figure 4.**
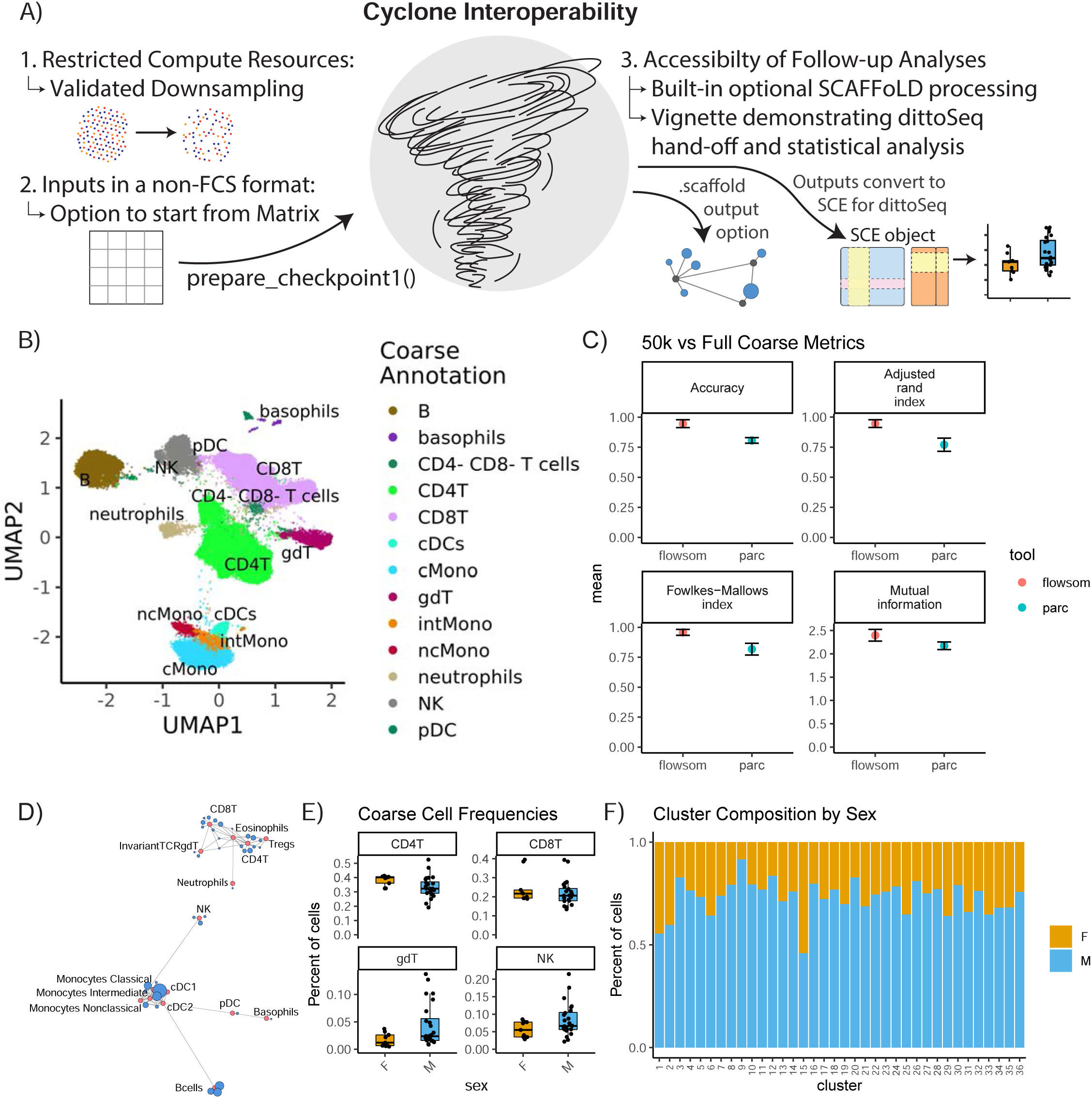
Accessibility of Cyclone - downsampling and interoperability with upstream and downstream processing. The dataset was down-sampled to 50k cells per sample and then run through cyclone. Clusters’ cell type identities were inferred by experts using cyclone plot outputs. Then, cyclone outputs from the full dataset were imported into R and read into a SingleCellExperiment object [cite, PMID in methods] so that further visualization and analysis could be carried out with dittoSeq. **A)** Overview of interoperability challenges and our solutions. **B)** UMAP from 50k down-sample run, colored by coarse annotations of one expert annotator. **C)** Comparison of per-cell annotations between the 50k down-sample versus the full dataset. **D)** Example Scaffold map export. **E)** Box plot showing per-sample cluster frequencies grouped by sex of the patients for for coarse-level cell types, created with dittoSeq’s dittoFreqPlot function. **F)** Stacked bar plot showing the percent of cells in each cluster by sex of the patients, created with dittoSeq’s dittoBarPlot function.

To optimize Cyclone interoperability with upstream and downstream processing, we considered batch correction as a primary upstream target, and SCAFFoLD map analysis and accessible visualization as primary downstream targets. For batch correction, we ensured a pathway exists to enter the Cyclone workflow with cytometry data in either a matrix format (as output by CyCombine (22)) or adjusted-FCS files (as output by CytoNorm (21)) to accommodate interoperability with outputs from both commonly used batch correction methods. CytoNorm’s adjusted-FCS files can enter into the pipeline as normal, but Cyclone’s prepare_checkpoint1() provides an entry point for CyCombine’s matrix format. The function takes the same primary inputs as a run without a batch correction step, except for accepting a matrix (with markers in rows and files in columns) instead of the FCS directory. The function then performs all the same optional steps (arcsinh transformation, control sample removal, and subsampling) before outputting a Checkpoint1.Rdata file. Afterwards, users can run Cyclone as normal to continue from step 2.

For powering SCAFFoLD map downstream analysis, we added an optional step directly into the cyclone pipeline; if landmark population FCS files are provided, an output *.scaffold file can be used to generate a SCAFFoLD map (15) via the “scaffold::scaffold.run()” command (**Figure 4D**). For powering accessible visualization and other downstream follow ups, we ensured that the colors of Cyclone plot outputs use color blindness accessible color palettes, and that data outputs can be used with dittoSeq--a color blindness friendly visualization tool (16). Although dittoSeq was designed for single-cell RNAseq data, it proves generalizable to other data modalities including high dimensional cytometry. To directly power dittoSeq integration and downstream statistical analyses, we provide an additional vignette showing how to 1) transform Cyclone data objects into a SingleCellExperiment ((27), Chapter 4) object compatible with dittoSeq, 2) generate useful visualizations, 3) add cell type annotations for each cluster, and 4) run statistics on differences in cluster or cell type frequencies between samples. Shown as examples are boxplots of how cluster frequencies compare between male and female subjects (**Figure 4E**) and the percent composition of each cluster in terms of subject sex (**Figure 4F**); these functions make probing metadata categories of interest easy to code and assist in producing publication ready visualization. These accessible (both in color palate and easily leveraged graphing functions) visualizations are readily created with dittoSeq from Cyclone outputs.

### The Cyclone pipeline generalizes to flow cytometry datasets

While optimization and development of Cyclone was initially for CyTOF datasets, we realized the need for such analysis techniques on a broader set of similar data modalities, including other cytometry platforms such as spectral flow cytometry. To evaluate Cyclone’s utility for this analysis, we applied the Cyclone pipeline to a spectral flow cytometry dataset of mouse liver samples from a transgenic mouse model for Hepatitis B Virus (HBV) response (28). Liver leukocytes collected from eight mice on day 8, the peak of the immune response in this model, were analyzed using a myeloid focused spectral flow cytometry panel (**Supplemental Table 2**). In a repeat experiment, day 8 liver leukocytes from nine mice were analyzed using a lymphoid focused panel (**Supplemental Table 3**). The Cyclone pipeline could be used as built, but we used config files to specify the co-factor for the arcsinh transformation to a value more typical of flow cytometry datasets (See “Spectral flow data generation and analysis – Cyclone analysis” in Methods). For both experiments, CD45+ live cells were provided to the Cyclone pipeline. After selecting the local minimum DBI, we identified and annotated 21 clusters in the T Cell panel (**Figure 5A**), and 17 clusters from the Myeloid panel (**Figure 5B**). Both panels were able to distinguish several T cell and NK cell subsets (**Supplemental Figure 4A**) as well as liver resident macrophages (Kupffer cells) and monocytes-derived macrophages (**Supplemental Figure 4B**). As was previously observed for CyTOF data, unsupervised clustering using Cyclone largely recapitulated the populations identified by manual expert gating at a coarse level (**Figure 5C, 5D**). It further enabled the identification of cell subsets across immune cell lineages (**Figure 5C, 5D**) which could have been missed by manual gating, such as NK cells subsets with various expression of KLRG1 or CD62L (**Figure 5A and Supplemental Figure 4A**). To evaluate the biological information contained within the unsupervised clustering, we took advantage of the presence of tetramer staining to identify the properties of the HBV-specific T cells. While the tetramer staining was not used as a clustering parameter, we found that the vast majority of CD8+ (**Figure 5E**) and CD4+ (**Figure 5F**) antigen-specific T cells were enriched in one or two effector T cell clusters with high expression of activation markers identified by Cyclone for each T cell subset (**Figure 5G**). Taken together, we observed that Cyclone performed well on spectral flow data and enabled the unsupervised identification of cell phenotypes that are associated with distinct biological features.

**Figure 5.**
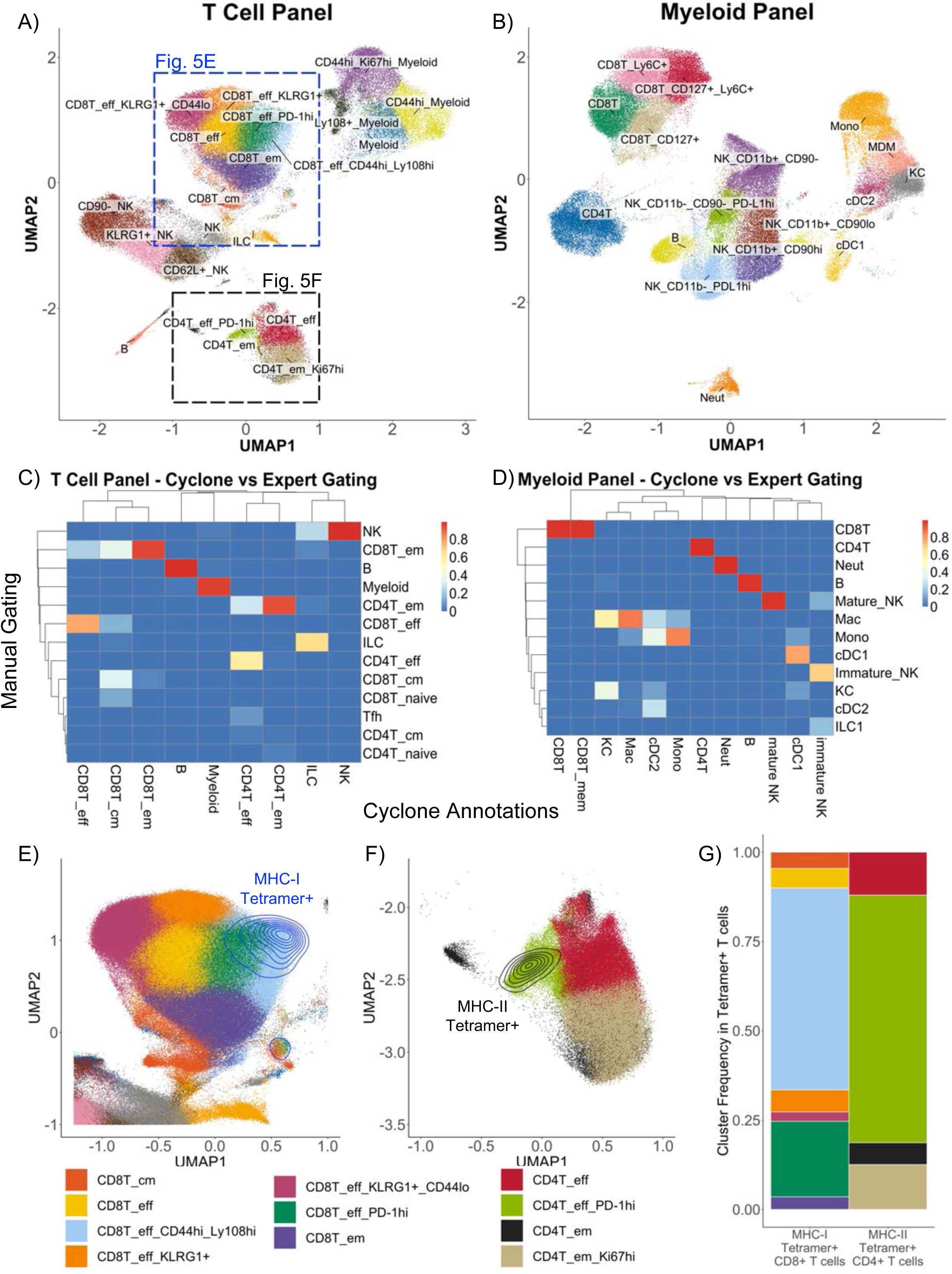
Utilization of Cyclone pipeline for spectral flow cytometry data. Liver leukocytes from HBVEnvRag−/− mice were harvested 8 days after adoptive transfer with wild type splenocytes for spectral flow cytometry analysis. Cyclone pipeline was run on this spectral flow cytometry data. **A)** “Fine”-level annotated UMAP of 22-color T cell-focused spectral flow cytometry panel (Supplemental Table 3) run on 9 mouse samples. **B)** “Fine”-level annotated UMAP of 25-color myeloid-focused spectral flow cytometry panel (Supplemental Table 2) run on 8 mouse samples. Three unidentifiable “junk” clusters were removed from this UMAP. **C)** Heatmap of “coarse”-level cell type annotations comparing expert manual gating identities (rows) to Cyclone cluster annotations (columns) in **A** (T cell-focused panel). **D)** Heatmap of “coarse”-level cell type annotations comparing expert manual gating identities (rows) to Cyclone cluster annotations (columns) in **B** (Myeloid-focused panel). **E)** Zoomed section of **A** showing density plot of HBV-specific MHC class I tetramer+ CD8+ T cells. **F)** Zoomed section of **A** showing expression of HBV-specific MHC class II tetramer+ on CD4+ T cells. **G)** Frequencies of “fine”-level cluster annotations among Tetramer+ CD8+ or CD4+ T cells. Tetramer+ cells were defined as events with fluorescence intensities 3 or more standard deviations above mean fluorescence in their respective channels.

### Cyclone enables unsupervised discovery of cell type compartmentalization within the tumor microenvironment

Understanding how the phenotype of individual cells relates to the function of multicellular compartments within tissues requires the ability to identify cellular phenotypes with multiple proteins while simultaneously quantifying the spatial distribution and interactions of these cells across large regions of tissue. Along with its applications demonstrated already, Cyclone provides a unique opportunity to analyze the spatial distribution of individual cell phenotypes as well as cellular neighborhoods in an unsupervised manner. To that end, we tested how the Cyclone pipeline compared to a prototypical imaging data analysis pipeline containing image visualization. First, we created a 7-plex immunofluorescent staining panel to be used on a colorectal tumor tissue (CRC1) consisting of a tumor marker (EPCAM), T cell markers (CD3, CD4, CD8) and myeloid markers (CD163, HLA-DR, XCR1) (**Figure 6A**). Next, we utilized DeepCell (29) segmentation software to demarcate individual cells by inputting nuclei (DAPI^+^) and membrane markers (CD3^+^, CD4^+^, CD8^+^, CD163^+^) and labeled each cell type annotation based on rational gating parameters (e.g., CD4^+^ T cell=CD3^+^CD4^+^CD8^−^XCR1^−^) (**Supplemental Figure 5A, table**). We found that determining the manual thresholding on certain markers such as CD163^+^ expression was not visually clear in designating a suitable cutoff as compared to CD3^+^ expression and that this could ultimately lead to variability in the frequency of cell types annotated (**Supplemental Figure 5A, histogram plots**). For example, threshold cutoff at 11 observed CD163^+^ expression on low background, segmented cells, while threshold cutoff 13 missed CD163^+^ cells. Thus, we opted for a middle ground by choosing threshold 12 for the downstream comparison with our Cyclone pipeline. To run the Cyclone pipeline, on this multiplexed immunofluorescence data we followed the following procedure. Once raw expression values of each marker for every cell have been curated and assigned to each individual cell identified by DeepCell, we obtained a cell per protein expression matrix that we can use to run the Cyclone pipeline (see methods). After DBI evaluation we chose grid size of 2×4, which has the lowest DBI value (**Figure 6B**). This resolution generated eight unique cluster spanning immune non immune cells. (**Figure 6C**). These clusters comprised of tumor cells expressing different levels of HLA-DR (Cluster 2 and 4), CD4^+^ and CD8^+^ T cells (Cluster 3 and 8), mononuclear phagocytes (MNPs) (Cluster 5, 6, and 7), and one cluster with low expression for all markers (Cluster 1) (**Figure 6D**). Interestingly, among the different MNPs cluster 7 is characterized by the high expression of XCR1^+^ and CD163^+^ and low HLA-DR expression. Since the combination of the Cyclone and DeepCell segmentation outputs provides these cluster identities (Cyclone) and x,y coordinates (DeepCell) for each cell within the same data frame, we leveraged this to evaluate the validity of where these XCR1+CD163^+^ cells can be located within the tissue. When overlaying Cluster 7 onto the image of CRC1, cells depicted as Cluster 7 (green arrows) we were able to confirm that they had both XCR1^+^ and CD163^+^ expression (**Figure 6E**). Additionally, these cells had an enriched spatial distribution within the stromal compartment as opposed to cDC1s and other clusters enriched in the tumor compartment (**Figure 6F, 6G, and Supplemental Figure 5B**). We, next also validated that XCR1^+^CD163^+^ cell types could be found in other colorectal (CRC2) and kidney (KID1) tumor samples subjected into Cyclone (**Supplemental Figure 5C**). However, these samples observed different stromal and tumor enrichments for CD4^+^ T cells, cDC1s, and XCR1^+^CD163^+^ cells compared to CRC1.

**Figure 6.**
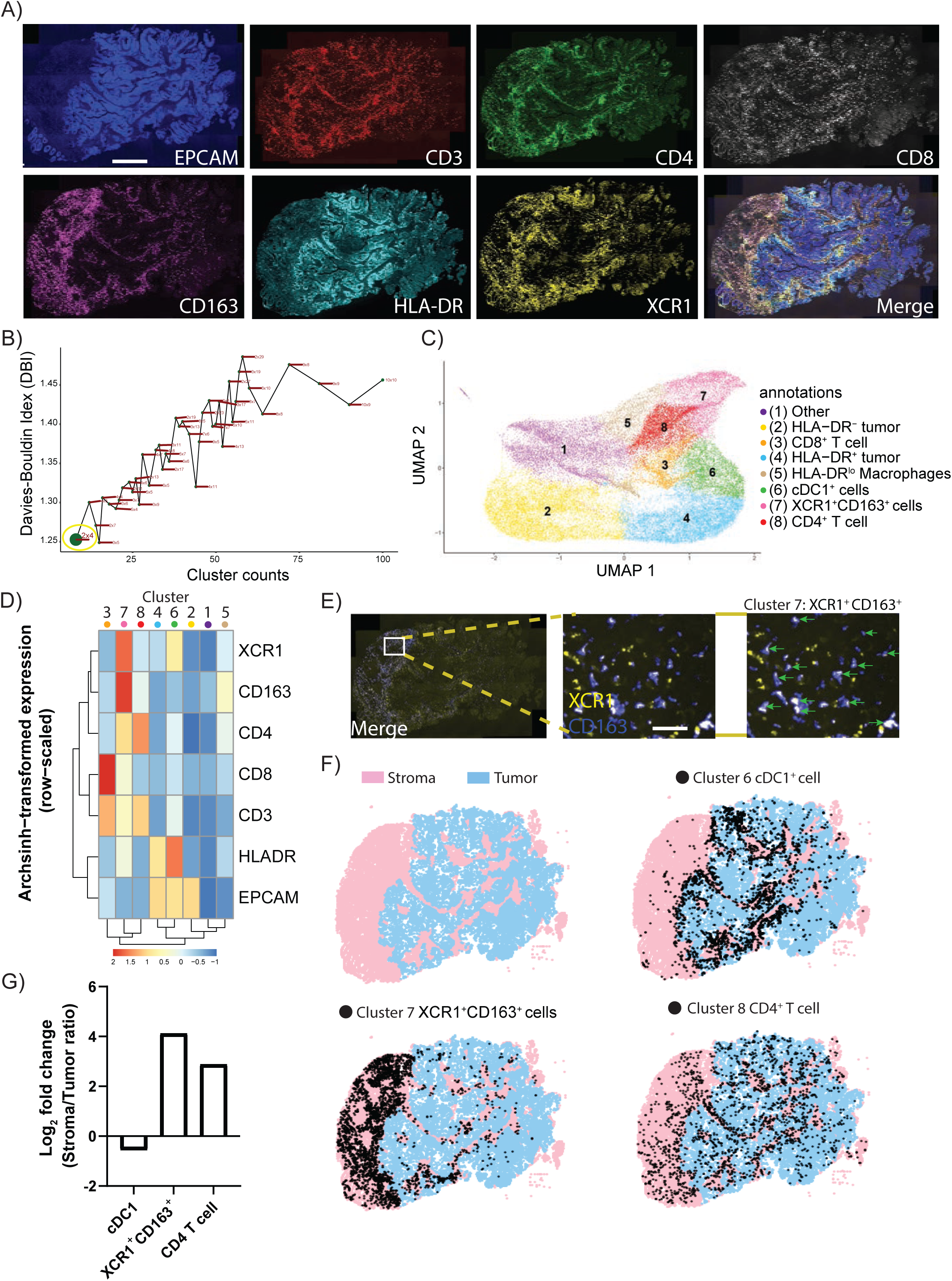
Utilization of cyclone pipeline for imaging data. **A)** Representative immunofluorescence imaging of colorectal tumor biopsy using tumor marker: EPCAM (blue), T cell markers: CD3 (red), CD4 (green), CD8 (white), and myeloid markers: CD163 (purple), HLA-DR (cyan), XCR1 (yellow) staining. Scale bar denotes 500μm. **B)** Scatterplot of the Davies-Bouldin index and cluster size over multiple iterations of Louvain clustering and varying parameters using 7 markers used in immunofluorescence as in A. Grid size 2×4 was chosen for downstream analysis (green dot, yellow circle). **C)** UMAP visualization of 45,177 cells from the colorectal tumor biopsy with specific populations annotated based on an arcsinh-transformed expression heatmap **(D)** of all markers row-scaled. **E)** Representative immunofluorescence imaging of tumor biopsy (merge; left) with inset (middle) of XCR1^+^ (yellow) and CD163^+^ (blue) staining. Cluster 7 from cyclone pipeline was overlaid and annotated (green arrows) representing XCR1^+^CD163^+^ cells. Scale bar denotes 20μm **F)** Full image representation (left) of stromal (pink) and tumor (blue) regions in tumor tissue biopsy (top left) with cDC1^+^ cell cluster 6 (top right), XCR1^+^CD163^+^ cell cluster 7 (bottom left), and CD4^+^ T cell cluster 8 (bottom right) overlays (black dots). **G)** Log2 fold change bar plot on stromal/tumor ratio of each cluster annotation in CRC1 tumor sample.

Notably, while conventional manual gating for XCR1^+^CD163^+^ cells is possible, the thresholding strategy of comparing independent markers with strict cutoffs likely miss out on what Cyclone identifies while leveraging the entire set of markers to identify cell types (**Supplemental Figure 5D**). Taken together, we highlight the successful application of the Cyclone pipeline on multiplexed imaging data in tumor tissue that could discriminate two different XCR1^+^ subsets and their spatial features within the tumor microenvironment.

## Discussion

In this work, we present the Cyclone pipeline—a versatile and accessible pipeline for performing, optimizing, and evaluating clustering on cytometry datasets. The pipeline takes in single-cell measurements, performs high-dimensional clustering, allows the user to select a clustering resolution with guided metrics, and provides outputs for facile cluster annotation and downstream analysis. We confirmed the fidelity of FlowSOM clustering to expert manual gating, as well as its performance on high-dimensional datasets: to date we have successfully applied the Cyclone pipeline with FlowSOM clustering to a 42-parameter CyTOF dataset of 50 million cells. We have released the pipeline code and documentation with the aim of making it accessible to the greater community.

Central to this work was to build a tool that would be easy to use by the research community, including wet-lab scientists, such that researchers are better empowered to engage with their cytometry datasets. In our collaborative research model, we brought together the computational rigor from data scientists with the perspectives, challenges, and biological intuition from wet-lab biologists for the design and development of the Cyclone pipeline. This included the consideration of high-performance computational resource access, which is not always available or accessible to wet-lab scientists. We therefore validated Cyclone’s performance on downsampled data to offer an option for running the pipeline locally on a personal laptop. This also included addressing the challenges of varied inputs (i.e., starting with varied file formats, either .fcs files or a more general matrix format), as well as computational hand-off to downstream analysis tools, either of which could erect a barrier to researchers less proficient with coding.

Cyclone is readily generalized to a variety of cytometry datasets beyond CyTOF data, demonstrating its versatility as a pipeline for high-dimensional datasets that can be extensible and adaptable as technologies continue to evolve. We demonstrate the application of Cyclone to spectral flow cytometry data, in this case in the setting of a mouse model of viral infection, as well as immunofluorescence imaging data of the tumor microenvironment. Excitingly, Cyclone has been further applied to a CO-Detection by indEXing (CODEX) (30, 31) dataset as well as a series of other CyTOF datasets (30) and spectral flow datasets (data not shown), further confirming its versatility.

These additional applications of Cyclone not only validated that the pipeline was functional in these settings, but also demonstrated its strengths in identifying elements of the biological systems that may be overlooked with manual gating. Clustering of the tumor microenvironment imaging data revealed a CD163+XCR1+ cell subset that was enriched in the stroma region. It is now well accepted that XCR1 expression defines the classical dendritic celIs of type 1 (cDC1) across a wide range of organisms (32). It is therefore tempting to suggest that these cells may represent a population of CD163^+^ DC1 which has been previously described in human breast and lung cancer patients (33). However, in this study, these CD163^+^ DCs were defined as a discrete subset of DCs distinct from both cDC1s and cDC2s and had the ability to efficiently trigger CD103 expression in CD8^+^ T cells *in vitro*, but the expression of XCR1 was not measured. The origin of these XCR1+ CD163+ cells remains unclear and more work is warranted. Nevertheless, we anticipate that the flexibility and operability of the Cyclone pipeline will help address this question as well as aid in the investigation of these cells’ spatial relationship within the tumor microenvironment and better define their potential role in the tumor immune response. In the spectral flow dataset, HBV-tetramer staining could be integrated with the clustering of T cell subsets to better phenotype these antigen-specific cells. We found that the majority of tetramer+ CD4+ or CD8+ T cells clustered together in their respective compartments, indicating a shared phenotype. Unsurprisingly for this day 8 timepoint in the immune response, MHC Class I Tetramer+ CD8+ T cells and MHC Class II Tetramer+ CD4+ T cells fell primarily into clusters identified as effector T cells with high expression of markers of activation, including CD44 and PD-1, and markers of high proliferative capacity, including Ly108 and TCF-1. Notably, those two markers that have been previously identified in antigen specific CD8+ T cells during LCMV chronic infection and cancer and have been associated to a cycling T cell stage which precede exhaustion program upon chronic stimulation (34).

This work has several limitations which can be explored or developed in future work. The set of clustering algorithms we compared for selection was restricted to four popular algorithms that are prevalent in cytometry analysis and have been elsewhere benchmarked (35), rather than us doing a more exhaustive search *de novo*. In addition, though we selected FlowSOM as our default and fully optimized algorithm, CLARA and FlowSOM performed similarly in our comparisons; while the user has the option to select CLARA clustering, the DBI optimization only works with FlowSOM in our implementation. We additionally noted in our evaluation of the clustering that the selected resolutions sometimes failed to capture low-abundance populations as their own cluster, such as cDC1s, eosinophils or antibody secreted cells (ASC). This is unsurprising and a common challenge in global clustering approaches, which can be remedied either by (1) “over-clustering” the data (i.e. selected a higher resolution such that you may better capture lower abundance populations but larger populations are further partitioned and need to be subsquently merged back together into a single population) or (2) “subclustering” the data (e.g. taking only the myeloid subsets and clustering them with relevant markers such that only those cells and markers are partitioning the space. We also noted decreased accuracy clusters defined by markers with a continuum of expression such as CD45RA (e.g. naïve v. memory T cells); however, because the placement of this manual gate on a continuum is somewhat arbitrary, modest discrepancy between the manually-defined abundance and cluster abundance seems of low importance as long as each are applied consistently across samples of interest. In addition, while we found these algorithms to be robust to downsampling, we have found that FlowSOM and CLARA could not accommodate a larger CyTOF data set of ∼90 million cells, regardless of resource dedication. Further optimization of those algorithms is needed to be able to accommodate increasingly large cytometry datasets. Finally, while we have invested in interoperability and extensive documentation and vignettes for ease of use, Cyclone could be even more accessible to a non-coding user base with the development of a graphical user interface (GUI) such as an R-shiny application (https://shiny.rstudio.com) or as a workflow in web-based tools such as the UCSF Data Library (https://datalibrary.ucsf.edu/), the Chan Zuckerberg Initiative’s CELLxGENE (https://github.com/chanzuckerberg/cellxgene) or CellEngine (https://github.com/primitybio/cellengine-python-toolkit), which could be the subject of future efforts. In sum, Cyclone takes a next step forward in the optimization and democratization of cytometry-based analysis tools to further power biological discovery.

## Methods

### Mass cytometry data generation and preprocessing

#### Sample collection

Blood samples from patients were obtained under IRB #11-07994 protocol approved by the UCSF IRB, the IRB of record for the study. Written informed consent was obtained from all patients. De-identified healthy donor sample was obtained from Vitalant Research institute (San Francisco, CA). Blood was collected into sterile EDTA vacutainer tubes (VWR International) and processed within 24 hours of collection.

#### Sample preparation

Peripheral blood mononuclear cells (PBMC) were isolated using Ficoll-Paque Plus (GE-Healthcare) density gradient centrifugation, after isolations cells were aliquoted in 0.5×10^7^ cells per vial in cell freezing media (10% DMSO in FBS) and cryopreserved.

#### CyTOF panel and staining

Mass cytometry was performed as described (36) with modifications. Briefly, primary conjugates of mass cytometry antibodies were prepared using the MaxPAR antibody conjugation kit (Fluidigm, South San Francisco, CA, USA) according to the manufacturer’s recommended protocol. Following labeling, antibodies were diluted in Candor PBS Antibody Stabilization solution (Candor Bioscience GmbH, Wangen, Germany) supplemented with 0.02% NaN3 to between 0.1 and 0.3 mg/mL and stored long-term at 4°C. Each antibody clone was titrated to optimal staining concentrations using unstimulated or anti-CD3/CD28 stimulated PBMC samples. All mass cytometry antibodies and concentrations used for analysis can be found in **Supplemental Table 1**. Mass cytometry experiments were performed over the course of nine separate experiments. Each PBMC sample was thawed at 37°C and washed in pre-warmed RPMI-1640 media (Sigma-Aldrich Life Sciences, USA) supplemented with 10% FBS (Gibco, ThermoFisher Scientific) in the presence of 250U Pierce Universal Nuclease for Cell Lysis (ThermoFisher Scientific, Rockford, IL), cells counted on Beckman Vi-Cell XR Cell Counter. Only samples with viability >75% were used (85% viability on average). 2.5×10^6^ cells/sample were stained for one minute with 25 mM Cisplatin (Sigma-Aldrich) in phosphate buffered saline (PBS) plus EDTA, before quenching 1:1 with PBS/EDTA/BSA to determine viability. Staining was performed on a shaker (90rpm). For staining, cells were first resuspended in cell staining media (CSM) (Fluidigm, South San Francisco, CA, USA) with 5ml Human TruStain FcX^TM^ block (Biolegend) for 5 min at room temperature to block Fc receptors, followed by staining with CXCR5 antibody in CSM (3mg/mL) for 30 min at 4°C. Cells were washed, fixed with Fix I Buffer from The Cell-ID™ 20-Plex Pd Barcoding Kit following manufacturer’s instructions (Fluidigm, South San Francisco, CA, USA) and barcoded by mass-tag labeling with distinct combinations of stable Pd isotopes diluted in Maxpar Barcode Perm Buffer (Fluidigm, South San Francisco, CA, USA) as described previously (5). Twenty barcoded samples were pooled into a single FACS tube (BD Biosciences) and stained with a cocktail containing surface markers antibodies (**Supplemental Table 1**) in a final volume of 1000ml CSM for 30 min at room temperature. Samples drawn at different timepoints per patient were barcoded together. Cells were then permeabilized with perm wash buffer (eBioscience, ThermoFisher Scientific) following manufacturer’s instructions and then incubated with a cocktail containing intracellular marker (**Supplemental Table 1**) antibodies diluted in perm wash buffer (eBioscience, ThermoFisher Scientific) for one hour at 4°C. Cells were finally stained with 191/193Ir DNA intercalator (Fluidigm, South San Francisco, CA, USA) diluted in PBC with 1.6% PFA (Electron Microscopy Sciences, Hatfield, PA, USA) 24h prior to data acquisition.

#### Data acquisition

For acquisition, cells were washed and resuspended at 1M/mL in deionized water + 10%EQ four element calibration beads (Fluidigm) and run on a Fluidigm CyTOF2 Helios mass Cytometer within one week of staining.

#### Data preprocessing (premessa -> bead normalization)

After data collection, we used the premessa pipeline (https://github.com/ParkerICI/premessa) to normalize data and deconvolute individual samples. From the individual sample files, normalization beads were excluded based on Ce140 and Eu153 signal. Single cell events were identified based on Ir191 DNA signal measured against event length, and CD45^−^ Pt195^+^ dead cells were excluded (**Supplemental Figure 1**). Potential batch effects were minimized by including a control sample from the same individual in each experimental run.

#### FCS modifications (addition of unique cell ids for tracking)

The FCS files do not contain cell identifiers. To accurately compare cell identity from either the manually gated annotations or the FlowSOM or PARC clusters annotated by two immunology experts, we added unique identifiers to all CD45+ live gated cells across all FCS files, and used these files for all downstream analyses. We created the unique identifiers by combined the sample identifier and the index of the single cells in each FCS file to generate unique identifiers, such as “<sample_id>_<cell_id>”. In this way, each cell gained a unique barcode id used for future comparative analyses.

#### Batch correction

The samples were processed across 8 batches. While the bead normalization of CyTOF data controls for the batch effects introduced due to instrument change, it does not address all factors affecting batch-to-batch variations (37). To evaluate the batch-effects, we first clustered the single cells using CLARA (23) and compared the batch compositions across clusters. We observed uneven distribution of batches across clusters (data not shown). To account for this residual batch-to-batch variation, we corrected the signal for batches using CytoNorm (21). We used the control samples (the same sample that was replicated across batches) to train the model and adjusted the batch-effects in non-control samples using nCells = 4k, nClus = 10 and the grid size of 5×5 (xdim=5; ydim=5). We determined that CytoNorm adjustment removed majority of batch-effects from the data (**Figure 3A**).

#### Manual gating

Main mass cytometry gating scheme can be found in **Supplemental Figure 1**, and shows exclusion of beads, debris, dead cells, CD45^−^ cells and granulocytes. Following this removal, we show the main gating strategy for identifying major immune cell populations from the mass cytometry dataset.

#### Generation of subsets

Different numbers of cells were selected from batch-corrected FCS files to produce specific subsets of the data: 1k subset = 1,000 cells per FCS file, 10k subset = 10,000 cells per FCS file, 50k subset = 50,000 cells per FCS file, Full subset = all cells from FCS files.

### Evaluation of runtime and memory usage for different clustering tools

To scope clustering algorithms for our pipeline, we tested four Python or R based widely used clustering tools for cytometry data: CLARA, FlowSOM, Phenograph, and PARC. CyTOF analysis allows users to capture hundreds of thousands of cells, and clustering such large datasets requires runtime- and memory-efficient tools that do not compromise clustering performance. One aim of our work was to design a scalable pipeline for large datasets (containing many samples, each with hundreds of thousands of cells). Therefore, we decided to compare the runtime and memory usage of the selected clustering tools. Different parameters affect the runtime and memory usage in different tools. We observed the parameters affecting the number of clusters were the ones that controlled algorithm runtime. To perform a fair comparison between the tools, we identified these parameters affecting cluster counts and used values that produced similar number of clusters across all four tools. We used k = 24 in CLARA; xdim=6 and ydim=6 in FlowSOM; k = 25 in PhenoGraph; and resolution = 1.3 in PARC. These parameters resulted in 30-37 clusters called by each tool. We measured the time for clustering and the memory usage on CentOS nodes of a high-performance computer cluster.

### Pipeline

#### Inputs

In order to begin a Cyclone run, metadata files “file_metadata.csv” and “marker_metadata.csv” and “config.yml” must be generated. These files are unique to your dataset, but Cyclone requires certain metadata to locate the FCS files and associate metadata with them. In the file metadata csv, column “file_name” records the name of each FCS file, “donor_id” denotes sample origin, “pool_id” denotes batch identification (if any), and “control_sample” is a Boolean indicating whether the sample is a control or not. The file metadata csv is created via any scripting language based on the FCS files present. To create the “marker_metadata.csv”, we provide “cyclone/make_marker_metadata_csv.R” which will read in an FCS file, and created a file with the following columns: channel_name, marker_name, used_for_UMAP, used_for_clustering, used_for_scaffold. It may be advantageous to use a marker for clustering, but not use that marker for UMAP calculation. Thus, the Boolean values for each marker in the “used_for_*” columns provided granular controls for using markers for UMAP, clustering, and scaffold analysis. The final file “config.yml” controls how Cyclone will be run. In brief, it contains the absolute path of FCS files, save location for pipeline outputs, location of metadata files, and parameter associated with data processing, UMAP, and clustering. Cyclone is able to cluster with both FlowSOM or CLARA, and the choice is specified in “config.yml”

#### Processing steps

At the start of the Cyclone run, the pipeline references the metadata.csv files to read in FCS data files and create a raw expression matrix. After arcsinh transformation of count values, a transformed expression matrix is created using values specified in config.yml. Cyclone also creates a “cell_metadata” object, which associates each cell/event with file and marker metadata. Next, Cyclone uses the uwot package with default parameters to calculate UMAP on the transformed matrix with the “used_for_UMAP” column of markers metadata file. Each cell/event UMAP dimensions are assigned to the cell_metadata object. Next, clustering is preformed using either FlowSOM or CLARA. In FlowSOM, a default series of grids (cyclone/grid_sizes.csv) is specified, and different resolutions and clustering parameters are calculated and evaluated with the Davies-Bouldin Index (DBI). After cluster optimization, the Cyclone pipeline exits to await user evaluation of the Cluster VS DBI plot (**Figure 1D, Supplemental Figure 2A**). If CLARA is used for clustering, the user must specify clustering parameters, and no DBI-based optimization is performed. After specifying a specific grid (FlowSOM) or specifying “k” in config.yml to calculate clusters (CLARA), clustering is calculated on the transformed matrix with the “used_for_clustering” column from the markers metadata file. Cluster assignment is stored in cell_metadata object. After clustering, Cyclone calculates cluster frequency matrices (raw and normalized) and calculates cluster median expression matrix. If scaffold analysis is selected, a gated directory of landmark FCS files is required. For each cluster, Cyclone gets the closest landmark population and stores this assignment in cluster_metadata. After calculating statistics, scaffold analysis is saved in a cluster_metadata object. With analysis completed, Cyclone outputs several helpful plots.

### CyTOF cluster annotation and benchmarking using manually gated populations

After clustering the data using FlowSOM or PARC, we performed manual annotation of the resulting clusters from both tools based on the median expression of markers. Since each individual may annotate clusters differently, we attempted to account for human-to-human variations in manual annotation of clusters by having two immunology experts to independently annotate the clusters from FlowSOM and PARC. We established “coarse” and “fine” levels of annotations. Coarse annotations describe cells of different compartment groups, (e.g. CD4+ T cells, CD8+ T cells, B cells, etc). Fine annotations further parse cell types into subtypes according to their phenotype (e.g., CD8+ T cells are split further into naïve, central memory, effector memory, and effector memory re-expressing CD45RA). We then calculated similarities between cluster annotations of single cells and the single cell annotation based on a third immunology expert gating of the CyTOF data (Ground Truth annotation). We performed this comparison for subsets of data multiple times and calculated averages. Specifically, we subsampled cluster annotation and manual gating annotation of randomly selected 10,000 cells and calculated accuracy, adjusted rand index, Fowlkes-Mallows index and mutual information. We repeated this process 10 times with different random seeds and calculated mean across the iterations for each similarity metric. We performed this analysis for FlowSOM (**Figure 1H, Supplemental Figure 2C**) and PARC annotations (data not shown) from both immunology experts.

### dittoSeq visualizations

Cyclone outputs (checkpoint1.Rdata and checkpoint8.Rdata) were used to create a SingleCellExperiment object (27) in R containing the arcsinh transformed expression matrix, umap embeddings, clustering, and cell and sample-level metadata. dittoSeq (16) functions dittoFreqPlot and dittoBarPlot were then used to create boxplots of cluster frequencies per sample and stacked bar plots of batch composition per cluster, respectively.

### Spectral flow data generation and analysis

#### Mice

WT C57BL/6 mice were purchased from Jackson Laboratory. HBVEnvRag^−/−^ mice were previously described (28). Briefly, HBVEnvRag^−/−^ mice were generated using HBV-Env^+^ mice [lineage 107-5D; gift from F. Chisari, Scripps Research Institute (38)] backcrossed to *Rag1*^−/−^ C57BL/6 mice for 15 generations. HBVEnvRag^−/−^ mice contain the entire envelope (subtype ayw) protein-coding region under the constitutive transcriptional control of the mouse albumin promoter. Young (3 weeks old, before weaning) or adult (8 to 12 weeks old) HBVEnvRag^−/−^ mice were given 10^8^ syngeneic splenocytes pooled from adult (8 to 12 weeks) WT mice in 0.5 ml of phosphate buffered saline via tail vein injection. Mice were maintained at the Laboratory Animal Resource Center (LARC) facility at UCSF where health and well-being was monitored daily by LARC staff. Experimental procedures were performed in accordance with IACUC-approved protocols and all efforts were made to minimize animal suffering.

#### Sample preparation

Mice were anesthetized in chambers with 1.5% oxygen and 3% isoflurane. Samples for the T cell-focused panel were isolated from the liver after perfusion and digestion. Briefly, mice were perfused via the inferior vena cava using digestion media [Hanks’ Balanced Salt Solution (HBSS), crude collagenase (0.2 mg/ml; Crescent Chemical), and DNase I (0.02 mg/ml; Roche Diagnostics)]. Livers were forced through a 70-μm filter using a syringe plunger, and debris was removed by centrifugation (30g for 3 min). Supernatants were collected and centrifuged for 10 min at 650g. Cells were isolated from the Percoll interface using a 60%:40% Percoll gradient. Samples for the myeloid-focused panel were isolated from the liver after 6 min of perfusion via the inferior vena cava using digestion media as above. Livers were chopped and further digested with liberase and DNase I (Roche Diagnostics) [1 Wünsch Units (WU) and 0.8 mg, respectively, in 10 ml of RPMI 1640 containing 5% FBS] for 30 min at 37°C in a shaking water bath. Livers were forced through a 70-μm filter, and debris was removed by centrifugation (30g for 3 min). Supernatants were collected and centrifuged for 10 min at 650g. Cells were isolated from the interface of a 60%:40% Percoll gradient.

#### Staining and acquisition

Cells were prepared as above. Samples stained with the T-cell focused panel were first stained with custom HBV-specific tetramers developed with the NIH Tetramer Core Facility at Emory University. Cells were stained first with MHC Class II Tetramer for 1 hour at 37°C protected from light. Next, these cells were then stained with MHC Class I Tetramer for 1 hour at 4°C protected from light. For both panels, cells were then stained with Live/Dead Fixable Blue (Thermo Fisher Scientific) according to manufacturer’s instructions. Next, surface markers on cells were stained according to standard protocols with anti-mouse antibodies detailed in **Supplemental Table 2** (Myeloid-focused Panel) or **Supplemental Table 3** (T cell-focused Panel). Finally, cells were fixed and permeabilized using FoxP3/Transcription Factor Staining Buffer Set (Thermo Fisher Scientific Cat# 00-5523-00) and stained with anti-mouse antibodies targeting intracellular markers according to standard protocols.

#### Data acquisition and preprocessing

Single color reference controls were collected for live unmixing with calculated autofluorescence immediately before fully stained sample acquisition. For sample acquisition, cells were run the same day as preparation and staining on an Aurora flow cytometer (Cytek) with 5-laser setup at the UCSF Flow CoLab. Following sample collection, spectral signatures were checked to ensure reliable unmixing and channel spillover was adjusted in SpectroFlo (Cytek). Prior to Cyclone and manual gating analyses, fcs files were gated on leukocyte size/granularity, singlets, live, and CD45+ events in FlowJo (**Supplemental Figure 4C**).

#### Cyclone analysis

Cyclone pipeline was run as described above, specifying an arcsinh transformation cofactor of 6000. For both panels, forward scatter, side scatter, CD45, Live/Dead Blue, and autofluorescence were excluded for UMAP generation and clustering. In addition, for the T cell-focused panel, channels for tetramers were excluded for UMAP generation and clustering. For the T cell panel, we identified 21 clusters, all of which were identified as specific lymphocyte populations (B cells, NK cells, T cells, or ILCs) or were assumed to belong to the myeloid compartment (**Supplemental Figure 4A**). For the myeloid panel, we identified 20 clusters. Among these clusters three were unidentifiable by markers in the panel (**Supplemental Figure 4B**) and excluded from subsequent analysis (**Figure 5B, 5D**).

#### Manual gating

Expert manual gating was performed in FlowJo to assign unique cell type identities to events. Example plots for T-cell focused gating strategy (**Supplemental Figure 4D**) and Myeloid-focused panel (**Supplemental Figure 4E**) are provided.

#### Annotation Comparison

FCS files for manual gates were exported from FlowJo then read into a flowFrame object in R using FlowCore package. Raw data was used to uniquely match events and assign annotations from manual gating and Cyclone clustering. Cells that received both manual gating and Cyclone clustering annotations were used to generate annotation concordance heatmaps (**Figure 5C, 5D**).

#### Tetramer+ Cell Visualization

Tetramer+ events were identified as having raw fluorescent intensities 3 or more standard deviations above mean intensity.

### Imaging data generation and analysis

#### Sample preparation

All patients consented by the UCSF IPI clinical coordinator group for tissue collection under a UCSF IRB approved protocol (UCSF IRB#20-31740). Samples were obtained after surgical excision with biopsies taken by pathology assistants to confirm the presence of tumor cells. Freshly resected samples were placed in ice-cold PBS or Leibovitz’s L-15 medium in a 50ml conical tube and immediately transported to the laboratory for sample labeling and formalin fixed for imaging analysis. Clinical data on three samples are denoted as follows: CRC1=IPICRC072, CRC2=IPICRC057, KID1=IPIKID090.

#### Staining and Imaging Immunofluorescent (IF) 7-plex panel

7-plex IF panel was created using the Ventana BenchMark Ultra (Roche Diagnostics) automated staining platform. All reagents were from Discovery (Ventana Medical Systems) and used according to manufacturer’s instructions, except as noted. Heat Induced Epitope Retrieval (HIER) was performed with the Cell Conditioning 1 (CC1) solution (cat. 950-124) for 64 min at 97°C. Primary antibodies used were CD3 (1:100, clone: D7A6E from Cell Signaling Technology), CD4 (RTU, clone: SP35 from Ventana), CD8 (1:100, clone: D8A8Y from Cell Signaling Technology), CD163 (1:250, clone: EPR19518 from Abcam), HLA-DR (1:500, clone: EPR3692 from Abcam), XCR1 (1:40, clone: D2F8T from Cell Signaling Technology) and EpCAM (1:50, clone: D9S3P, from Cell Signaling technology). The tissue was counterstained with DAPI (Akoya cat. FP1490) for nucleus localization. The staining was conducted in two cycles: first cycle had CD3, CD4, CD8, CD163, HLA-DR and XCR1 and the second cycle had EpCAM. Both cycles had DAPI. The slide was scanned using a whole slide scanner after each staining cycle. Finally, the images from both cycles were registered to achieve the 7-plex image shown in Figure 6.

#### Data preprocessing

Cell segmentation was performed by utilizing ark-analysis (v0.2.9) DeepCell (29) software with nuclei (DAPI) and membrane (CD3, CD4, CD8, CD163) as modalities for segmentation. Mean fluorescent intensity was measured from each cell ROI and then arcsinh transformed to input into Cyclone pipeline as “trans_exp”.csv file. Manual gating was performed in a custom Napari application version 0.4.14 to classify cells as positive or negative for each marker. For stroma versus tumor region separation, Qupath (version 0.3) software was used to annotate each region using EpCAM for colorectal or PanCK for kidney as a marker reference. Finally, data cluster labels generated from Cyclone pipeline or manual gating were generated as a csv file corresponding to each cell ROI and integrated with segmented imaging file for cluster overlay in Napari.

#### Cyclone analysis

After determining an optimal grid size, the Cyclone output heatmap showing arcsinh transformed and scaled marker expression data was used to annotate clusters and scaled by row. Log2 fold change of each cluster in stroma versus tumor was determined by calculating the frequency of stromal over frequency of tumor in each cluster and transformed by log of 2. Contour plots comparing Cyclone versus manual gating was generated in python (version 3.8.12) by plotting arcsinh transformed cell ROIs annotated as XCR1 and CD163.

## AUTHOR CONTRIBUTIONS

RKP and RGJ contributed equally to this manuscript. RKP, RGJ, IK, NDC, TC, NWC, BS, DB, LLJ, JMJ, JP and LA, generated, and analyzed data, and contributed to the manuscript by providing figures, tables and important intellectual contributions. RKP, RGJ, AJC and GKF had full access to all the data in the study and take responsibility for the integrity of the data and the accuracy of the analyses. RKP, RGJ, IK, DB, NDC, AJC and GKF wrote and edited the manuscript. RKP, RGJ, BS, DB, AAR and LLJ performed computational analyses of the data. MN and NC were part of the UCSF Immunoprofiler team who performed tissue preparation and staining, for the immunofluorescence performed on the different human tumor specimens. RKP, RGJ, BS, DB, AAR developed the Cyclone package and wrote the tutorial. AE and IK performed computational analysis of the multiplexed immunofluorescence data on human tumors. MFK, SC, JD AJC and GKF were actively involved the direction of projects and provided financial resources for the obtention of the data. AJC and GKF co-led this project. All authors edited and critically revised the manuscript for important intellectual content and gave final approval for the version to be published.

## ACKNOWLEDGEMENTS

We thank all members of the Combes and Fragiadakis labs, UCSF Bakar ImmunoX Initiative, UCSF CoLabs and the UCSF Immunoprofiler Consortium for discussion and guidance while developing this study. We would like to thank Drs David Erle, Michael G Kattah, and Elvira Mennillo for scientific discussion and editing the manuscript. We would like to particularly thank Isabelle Tingin, Garry Shumakher, Elizabeth Edmiston and Meghan Zubradt for their constant support and help during this study. Acquisition and analysis of certain human samples described in this study was partially funded by contributions from AbbVie, Amgen, Bristol-Myers Squibb, and Pfizer as part of the UCSF Immunoprofiler Initiative. Data acquisition was further supported by R01DK103735 and P30DK026743. Further support came from the Bakar ImmunoX Initative and ImmunoX Computational Biology Initiative at UCSF funding for AJC, GKF, DB, RKP, and RGJ. Finally, we thank all patients and their families, for placing their trust in us.

---

**Supplemental Figure 1.**
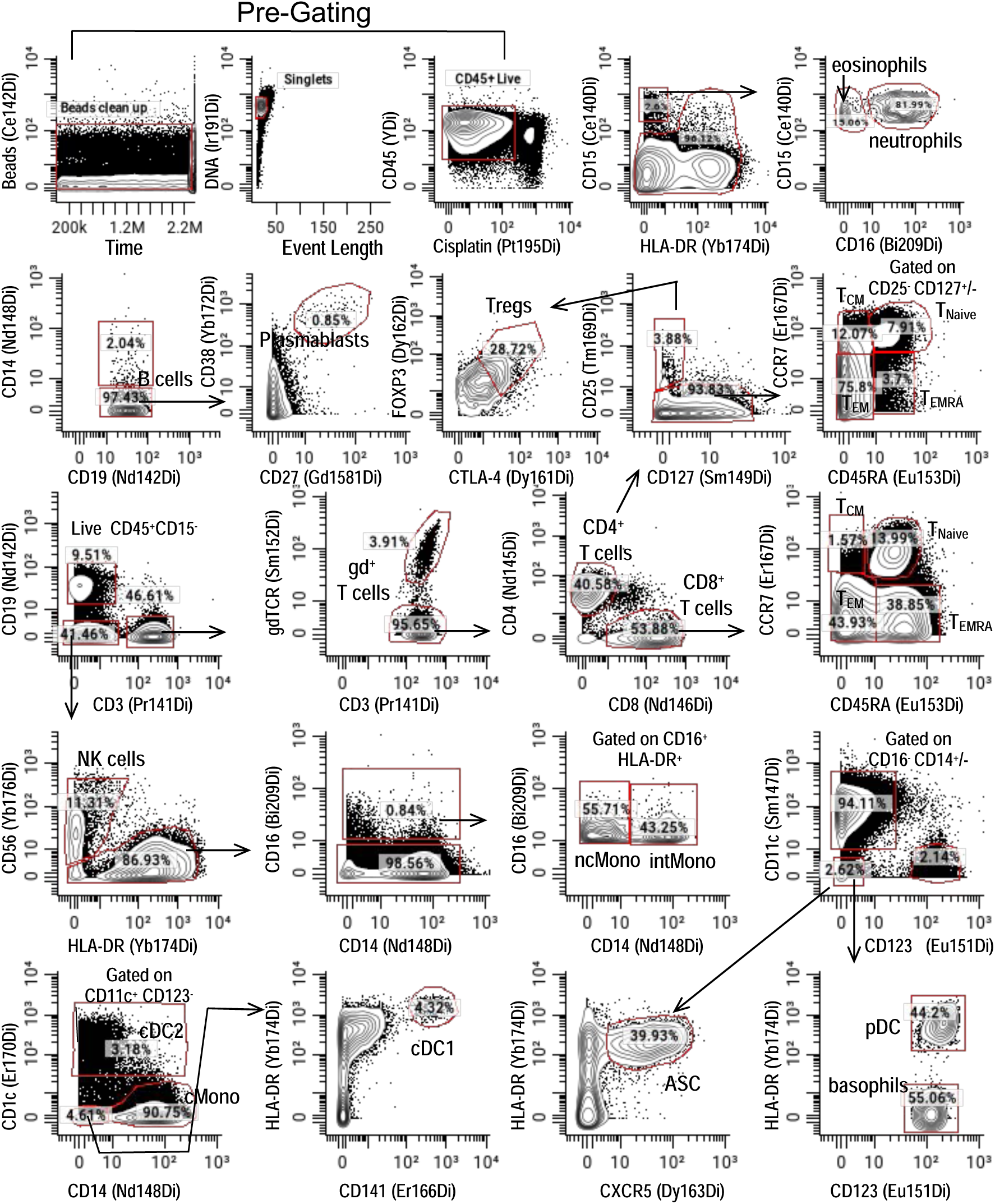
CyTOF manual gating of human PBMC. **A)** Pre-Gating: gating out beads, debris, dead cells, RBC and granulocytes. **B)** Hierarchical gating was applied to identify 22 “landmark” immune populations: CD14^+^ CD16^−^ classical monocytes, CD14^−^CD16^+^ nonclassical monocytes, CD14^+^ CD16^+^ intermediate monocytes, cDC1, CD2, pDC, basophils, Natural Killer cells, regulatory CD4^+^ T cells, CD4^+^ T cells (Naive, T_CM_, T_EM_, T_EMRA_), CD8^+^ T cells (Naive, T_CM_, T_EM_, T_EMRA_), γδ^+^ T cells, B cells, ASC (antibody producing cells), plasmablasts. Not shown gating of eosinophils (CD15^+^ CD16^+^ HLA-DR^−^) and neutrophils (CD15^+^ CD16^−^ HLA-DR^−^).

**Supplemental Figure 2.**
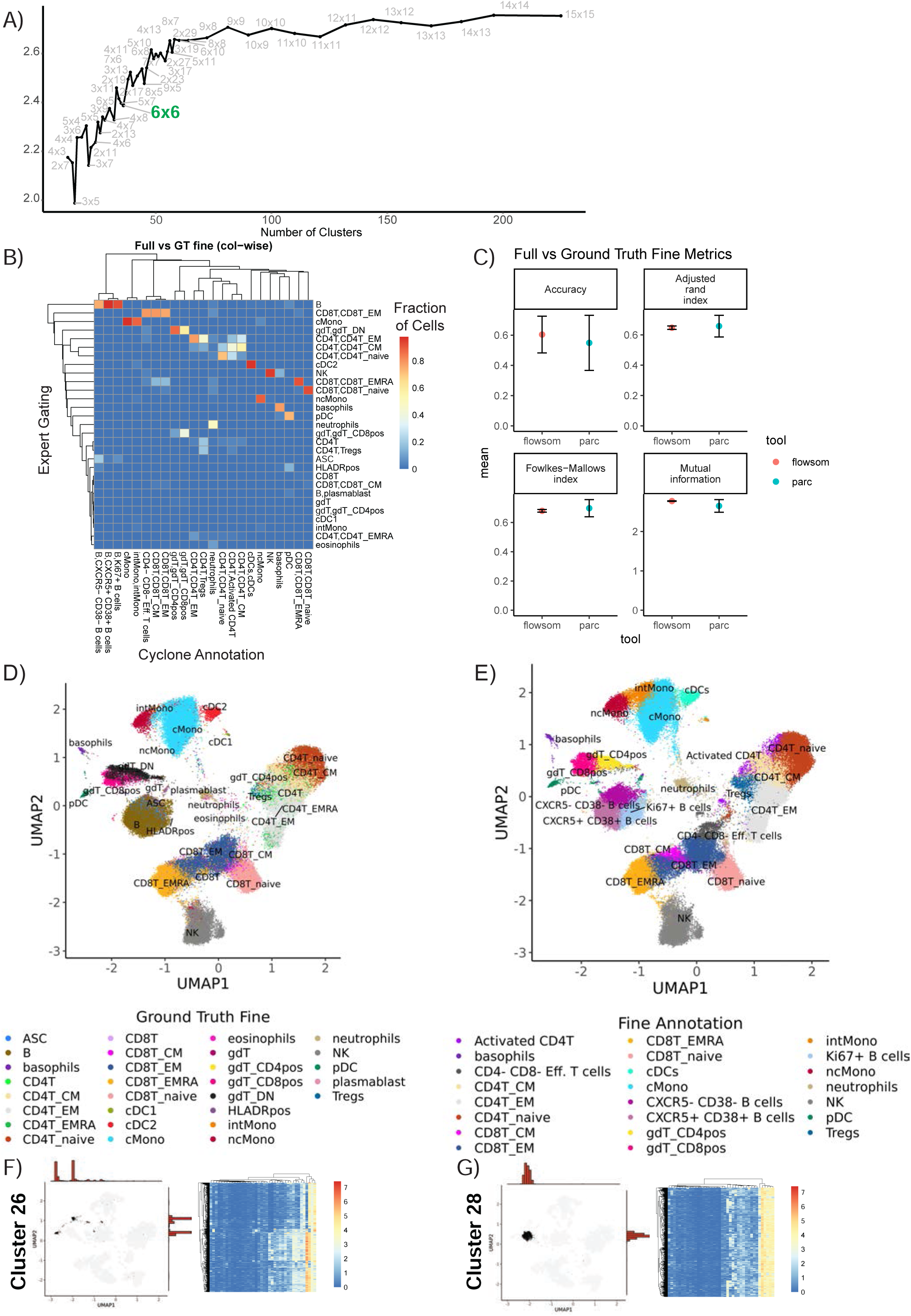
Assessment of “Fine”-level annotations and metrics for evaluating cyclone outputs. **A)** Full Davis-Bouldin index plot showing up to 200 potential clusters identified through FlowSOM. **B)** Heatmap of full dataset “fine”-level annotations identifying cell types and cell subtypes, based on ground truth (GT) manual gating (rows) compared to annotated FlowSOM clusters (columns). **C)** Comparison metrics based on “fine”-level annotations from two individuals. Various performance metrics were used to assess the accuracy of clusters called in the FlowSOM clustering compared to ground truth. **D)** Ground Truth expert cluster “fine”-level annotation identifying broad cell types and specific cell subtypes based on manual gating. **E)** FlowSOM clustering “fine”-level annotations based on CyTOF panel expression. **F)** Depiction of a cluster dispersed across UMAP space (Cluster 26) with a heterogenous protein expression profile compared to **G)** a cluster with uniform protein expression and tight UMAP localization (Cluster 28).

**Supplemental Figure 3.**
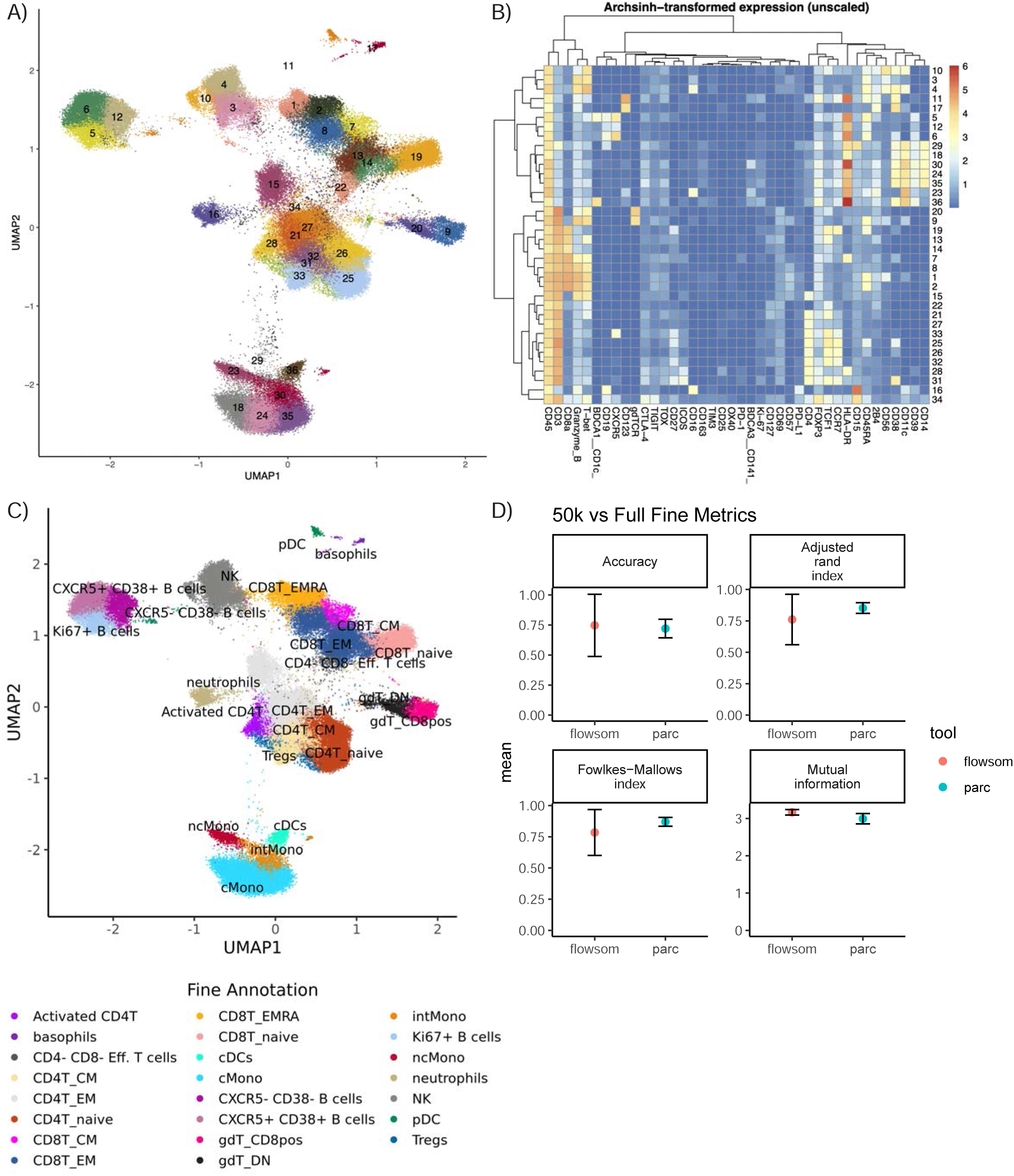
“Fine”-level annotations after running cyclone on the downsampled dataset. The dataset was down-sampled to 50k cells per sample and then run through cyclone. Clusters’ cell type identities were inferred by experts using cyclone plot outputs. **A)** UMAP annotated by cluster number. **B)** Heatmap of median archsinh-transformed expression (unscaled) per cluster, used to annotate clusters. **C)** UMAP from 50k down-sample run, colored by fine annotations. **D)** Comparison of per-cell annotations between the 50k down-sample versus the full dataset. Various performance metrics were used to assess the accuracy of clusters called in the downsampled dataset compared to the full dataset.

**Supplemental Figure 4.**
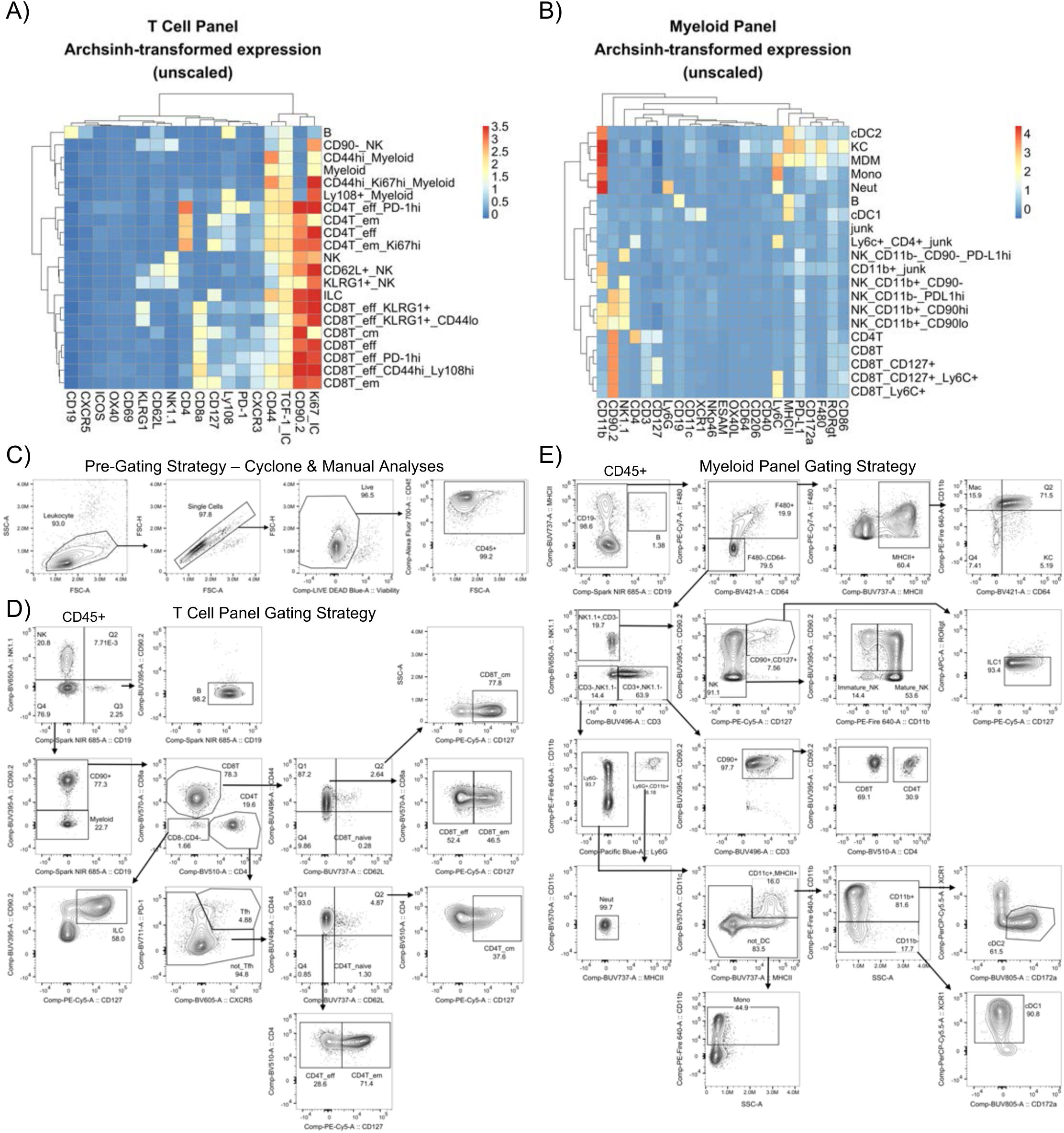
Spectral flow cytometry cell type identification. **A)** Heatmap of markers used for UMAP generation, clustering, and identification for “fine”-level cyclone clusters for the spectral flow cytometry dataset with a T cell-focused panel presented in Figure 5A. **B)** Heatmap of markers used for UMAP generation, clustering, and identification for “fine”-level cyclone clusters for the spectral flow cytometry dataset with a Myeloid-focused panel presented in Figure 5B. **C)** Representative two-dimensional flow plots demonstrating pre-gating on live CD45+ cells before analysis with either Cyclone or expert manual gating in FlowJo. **D)** Manual gating strategy for samples in the T cell-focused panel. **E)** Manual gating strategy for samples in the Myeloid-focused panel.

**Supplementary Figure 5.**
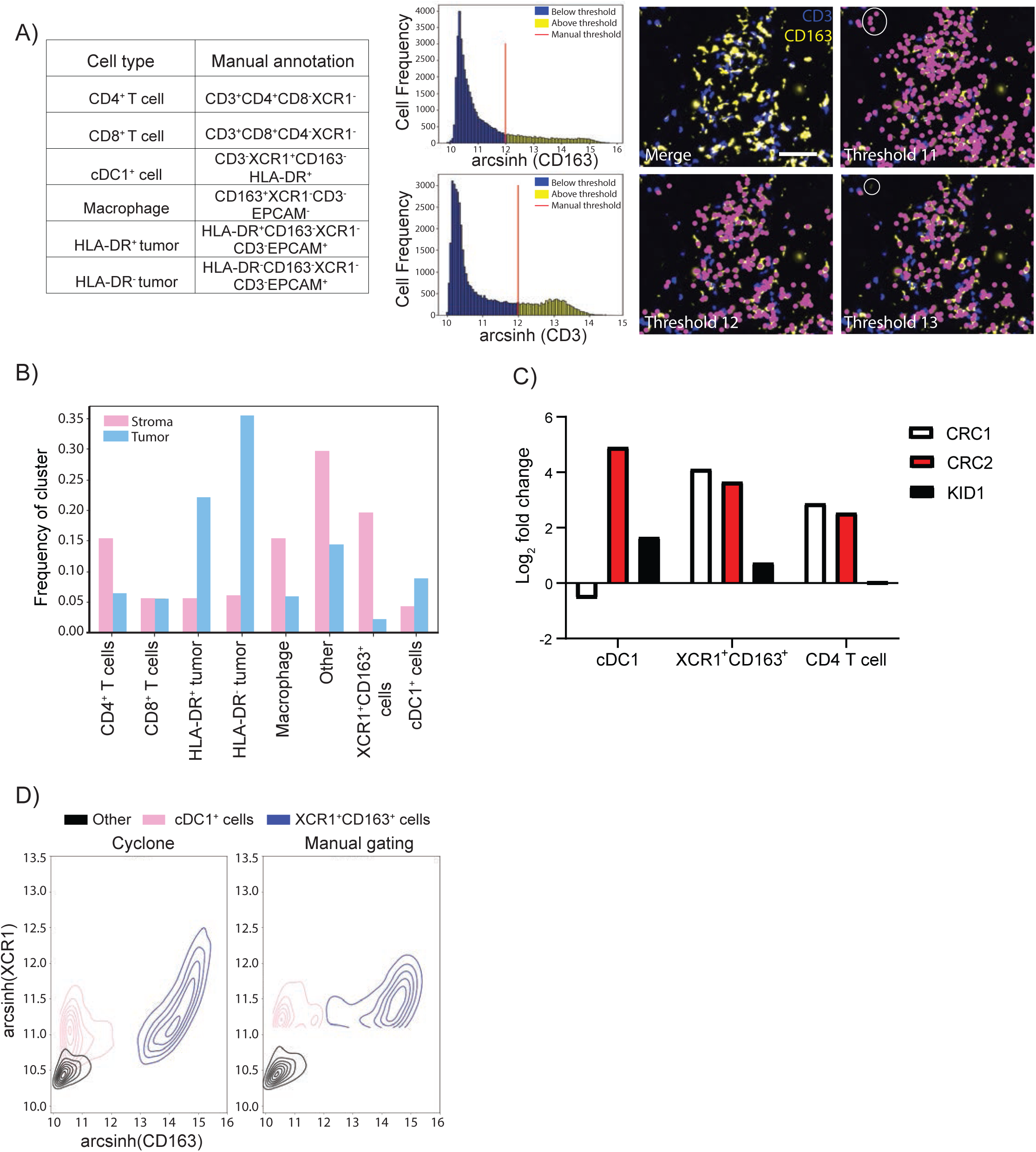
Resolution of manual gating and Cyclone clustering. **A)** Table (top, left) depicting cell type annotations based on the 7 markers used as in Figure 6A and cell frequency histograms with manual gating thresholds (red line) on above (yellow) or below (blue) threshold for CD163 and CD3 marker. Thresholds were set on CD163 and CD3 at 11, 12, and 13 to denote the cutoff and visualization of annotated macrophage^+^ cells (pink dots). Circled white regions indicate examples of over-thresholding (threshold 11) and under-thresholding (threshold 13). Scale bar denotes 20μm. **B)** Frequency bar plot of each cluster annotation divided into stromal (pink) or tumor (blue) compartments. **C)** Log2 fold change bar plot on stromal/tumor ratio for each annotated cluster in CRC1, CRC2, and KID1 tumor samples. **D)** Scatterplot of cDC1^+^ (pink), XCR1^+^CD163^+^ (blue) cells, and other between cyclone pipeline (left) and manual gating (right).

